# KPGminer: A tool for retrieving pathway genes from KEGG pathway database

**DOI:** 10.1101/416131

**Authors:** A. K. M. Azad

## Abstract

Pathway analysis is a very important aspect in computational systems biology as it serves as a crucial component in many computational pipelines. KEGG is one of the prominent databases that host pathway information associated with various organisms. In any pathway analysis pipelines, it is also important to collect and organize the pathway constituent genes for which a tool to automatically retrieve that would be a useful one to the practitioners. In this article, I present KPGminer, a tool that retrieves the constituent genes in KEGG pathways for various organisms and organizes that information suitable for many downstream pathway analysis pipelines. We exploited several KEGG web services using REST APIs, particularly GET and LIST methods to request for the information retrieval which is available for developers. Moreover, KPGminer can operate both for a particular pathway (single mode) or multiple pathways (batch mode). Next, we designed a crawler to extract necessary information from the response and generated outputs accordingly. KPGminer brings several key features including organism-specific and pathway-specific extraction of pathway genes from KEGG and always up-to-date information. Thus, we hope KPGminer can be a useful and effective tool to make downstream pathway analysis easier and faster. KPGminer is freely available for download from https://sourceforge.net/projects/kpgminer/.

## 1. Introduction

Biological pathway is defined as a collection of genes or proteins that are functionally related to each others to perform some biological activities such as signaling or regulatory activities. Some of the on-line pathway databases are KEGG [1], Reactome [2], Wikipathways [3] etc. where pathways related to signaling, metabolomic, cellular processes, diseases, genetic information are stored for various organism.

Pathway analysis is an important downstream component for many bioinformatics pipelines. One of the important aspects of a pathway analysis task set is to conduct enrichment test with already annotated pathways. This enrichment analysis include evaluating the enrichment of *de novo* gene sets (either computationally predicted or experimentally determined) with those already annotated pathways. Azad *et al.* designed a method called VToD [4] for identifying cancer-related gene modules, which were validated with known pathways from databases including KEGG [1] and GO terms [5] using gene set enrichment test. Another example of gene set enrichment analysis is to check the overlap of a particular set of interest e.g. differentially expressed genes with those annotated pathways. All of these enrichment tests require a set annotated pathways presented in the databases such as KEGG [1]. One of the sources to collect such annotated pathway sets is the Molecular Signatures Database (MSigDB) [6] which stores gene sets from well known pathway databases like KEGG [1] and Reactome [2]. But these gene sets are static and the pathway annotations are always updating. Hence, it is required to have a tool for retrieving up-to-date gene collections for users. Moreover, those collections aren’t organism-specific.

In this article, I present a standalone tool called *KPGminer* that retrieves the pathway genes from KEGG [1] for all the organisms every time it runs. This provide always up-to-date and organisms specific information which can be stored in the local machine for conducting downstream pathway analysis using some statistical methods such as hyper-geometric tests. I hope, this tool can be very useful for the researchers and contribute to their bioinformatics pipelines.

## 2. Implementation

Figure 1 shows the main KPGminer interface. When user opens KPGminer tool it loads all the available organisms in KEGG database by making HTTP web request via a web API call using REST protocol. This REST API protocol is available in the KEGG website for the developers’ use. Once, all the organisms are loaded successfully, the main interface of KPGminer popsup will a dropdown box populated with all the organism name. Next, user has to select a particular organism for the that list, another HTTP web request takes place for retrieving all the pathways currently available in KEGG database for that particular organism. The response of that request is then parsed to get the list of those pathways and a listbox gets populated with them. User can pick one or more pathways from that list which will be shown in another listbox (called selection listbox).

**Figure 1:**
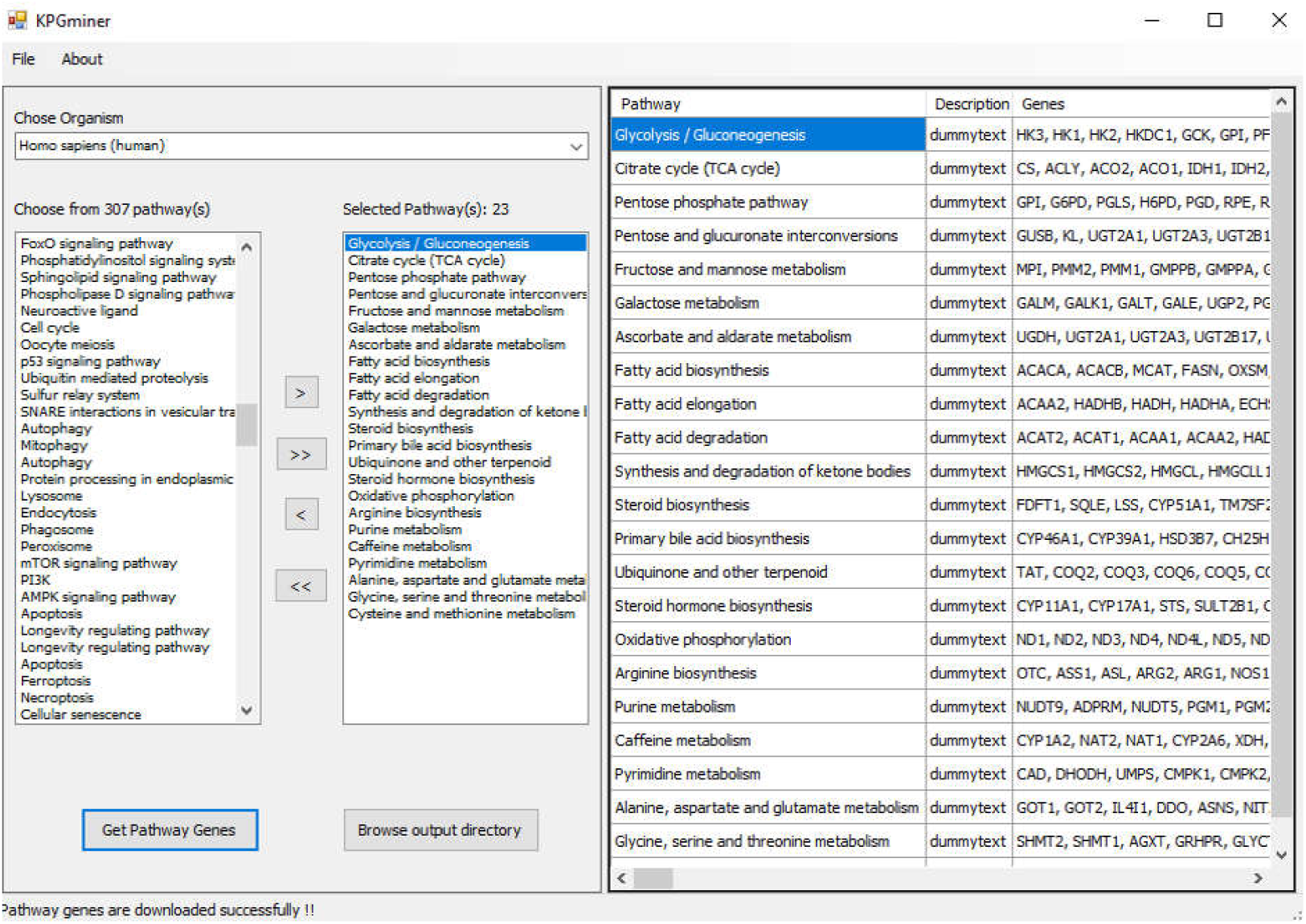
Main interface of KPGminer tool

To get all the pathway genes for those selected pathways, user press a button which makes another HTTP web request. KPGminer reports pathway genes in single or batch mode depending on the number of selected pathways. Once loaded all the pathway genes the results are shown in the right panel on the main KPGminer interface. Finally, to all of these pathway genes are can be saved in a file by clicking a button which asks a place to save that file in the local directory. The file is saved with a *.gmt* extension just as similar to the MSigDB for the convenience of users. A tooltip label keeps providing messages for every stages of KPGminer in retrieving pathway genes for selected pathway(s) for a particular organism. Table 1 shows the KPGminer metadata.

**Table 1:** KPGminer Metadata

|  | Technology used |
| --- | --- |
| Version | v1.0.0 |
| Language | C#.Net |
| Platform | Microsoft .Net platform |
| Operative systems | Windows |
| HTTP web request | REST API |
| Information retrieval technique | In-house built crawler |

## 3. Discussion

KPGminer has several useful features. First, even though KEGG provided necessary APIs for retrieving those information, a single platform to facilitate organism-specific and pathway-specific (single or batch mode) information retrieval may be advantageous for practitioners by abstracting their corresponding lower-level implementations. Second, while loading, KPGminer starts requesting KEGG databases for pathway information, which indicates that it always brings the up-to-date information. Third, KPGminer is a open source and free software that can help scientific communities to conduct pathway analysis required with KEGG pathway databases.

In this version of KGPminer, there is one limitation which is in batchmode (for multiple pathways) operation, it creates HTTP web request for each pathways separately, which is a time consuming. But this limitation can be overcome by exploiting multi-threading approach by making each HTTP web request running in a single thread, which can be implemented in future versions of KPGminer. In future I also hope to extend this tool for retrieving information from other pathway databases including Reactome [2], Wikipathways [3] or GO [5] database. I hope KPGminer can be a very useful tool for the researchers in their pathway analysis.

